# Metabolomics and microbiome reveal impacts of rhizosphere metabolites on alfalfa continuous cropping

**DOI:** 10.1101/2021.07.25.453728

**Authors:** Ruiting Wang, Jinxin Liu, Wanyi Jiang, Pingsheng Ji, Yonggang Li

## Abstract

Alfalfa long-term continuous cropping (CC) can pose a serious threat to alfalfa production. However, the mechanism of alfalfa CC obstacle is unclear as of today. In this study, we determined physic-chemical property, microbial population structure, and metabolite differences of alfalfa rhizosphere soils with CC for 1, 7, and 14-years based on analysis of metabolomics and microbiomics. Shifts of functional microorganisms in rhizosphere soil were analyzed, key metabolites and their effects on alfalfa seeds, seedlings and root rot pathogens were assessed. Based on anlysis, p-coumaric acid and ferulic acid on alfalfa seed and seedling growth and root rot pathogens were basically consistent with the influence of CC obstacles in the field. With the increase of CC years, the microbial community in soils changed from fungal to bacterial, and beneficial microorganisms decreased with the increase of CC years, which echoed the performance of alfalfa CC obstacles. The autotoxicity of p-coumaric acid was the strongest.This study fully proved that the continuous accumulation of autotoxic substances in alfalfa rhizosphere was the key factor causing alfalfa CC obstacles.

## Introduction

Alfalfa (*Medicago sativa* L.), also called lucerne or purple medic, is a perennial, clover-like, leguminous plant of the pea family (Fabaceae). It is widely grown primarily for hay, pasturage and silage in the United States, Europe and Asia (Graham & Vance, 2003) and is the main animal feed. The Ministry of Agriculture of the People’s Republic of China reported that cultivated area of alfalfa in China was 4.7 million ha with a production of 32.17 million tons from the end of 2015 to 2017. Among them, the commercial alfalfa planting area was 0.43 million ha, and the high-quality alfalfa planting area was 0.21 million ha with a production of 1.8 million tons. It was anticipated that the planting area of high-quality alfalfa in China will increase by 0.2 million ha by 2020 with a production of 3.6 million tons. However, with the continuous expansion of alfalfa planting area, alfalfa production capacity decreased year by year in fields continuously planted for more than 4 years. Reduced production capacity was reflected in that root activity declined gradually, seed germination rate decreased seriously, root rot incidence increased year by year, and it was difficult to survive resowing or replanting (Rong et al, 2016) This phenomenon is a typical continuous cropping (CC) obstacle of alfalfa.

There are two main explanations on the formation of CC obstacles of alfalfa in the literature. The first viewpoint is that alfalfa is a crop with deep root system and high water and fertilizer consumption. After continuous growth for many years, it will lead to soil water and fertilizer deficit, which causes a large area decline of alfalfa growth (Wu et al, 2015; Molero et al, 2019). Li et al. (2010) and Xia et al. (2015) also believed that CC of alfalfa led to deterioration of soil physical and chemical properties that are difficult to restore. However, other scholars hold different points of view. Li et al. (2010) found that planting alfalfa in the same land for more than 10 years could lead to serious land degradation and significant decline of alfalfa yield in the Loess Plateau. Viliana & Ognyan (2015) stated that certain numbers of roots would disappear after each harvest in the CC process of alfalfa, which could provide organic materials for the soil and increase soil organic matter contents. Jiang et al. (2007) reported that soil quality deteriorated but alfalfa yield increased in the first nine years, while soil quality tended to recover but alfalfa yield decreased in the following years. Our field studies also indicated that CC obstacle still occurred even if fertilizers and water were sufficient (Li, Y. G., unpublished). These studies demonstrated that soil moisture and fertility were not the primary factors causing the problems in alfalfa production, such as slow seed germination, delayed returning green, poor growth, yield and quality decline, and severe root rot in CC for more than 4 years.

Another explanation is that the continuous accumulation of autotoxic allelochemicals causes CC obstacle of alfalfa. Allelopathy is defined as the direct/indirect harmful/beneficial effect via the production of chemical compounds that escape into the environment (Rice, 1984). Allelopathy is a biological phenomenon by which an organism produces one or more biochemical substances that influence the germination, growth, survival, and reproduction of other organisms in the same community (Zuo et al, 2015). It was reported that alfalfa produced a number of useful phytochemicals, including soyasapogenol glycoside B (Wyman-Simpson et al, 1991 ), medicarpin and isoflavonoid (Miller, 1988), chlorogenic acid, ferulic acid, p-hydroxybenzoic acid, caffeic acid, coumarin, ferulic acid (Rong et al, 2016; Abdul-Rahman & Habib, 1989), amic acid, hydroxybenzoic acid, coumarin and tricinon (Zheng et al, 2018). These phenolic acids had inhibitory effects on plant seedlings, plant photosynthesis and respiration (John & Sarada, 2012; Li et al, 2012; Zhang et al, 2016; Biumal et al, 2019). Chung et al. (2000) reported that chlorogenic acid was involved in alfalfa autotoxicity. Rong et al. (2016) found that contents of coumarin, ferulic acid, chlorogenic acid and caffeic acid varied in 18 alfalfa varieties, with coumarin and chlorogenic acid being significantly higher than ferulic acid and caffeic acid. In other studies, phenolic acid such as p-coumaric acid inhibited photosynthesis and enzymatic activities of PG1, CG6PDH, AID and OPPP, which was detrimental to plant and root growth and altered morphological and physical structures of alfalfa roots (Zheng et al, 2018; Rong, 2017). These studies suggested that continuous accumulation of autotoxic substances in alfalfa rhizosphere might be the primary factor causing alfalfa CC obstacle; however, direct evidence supporting the theory is lacking.

Methods available for studying CC obstacle, include multi-omics, such as high throughput isolation (culturomics), analyzing structural and functional changes of plant rhizosphere microorganisms (microbiomics), targeting the taxonomic composition (metabarcoding), addressing the metabolic potential (metabarcoding of functional genes, metagenomics), and analyzing components of plant rhizosphere exudates (metabolomics). Through metabolomics, we can quickly understand the metabolic changes of organisms under the stimulation of different biological factors and environmental factors, search for biomarkers with the purpose of identifying metabolites related to various diseases and environmental exposure (Nicholson & Lindon, 2008, Liu et al, 2018). Through microbiomics, we can have a comprehensive understanding that plant surfaces and interior parts are populated by myriads of bacteria, fungi and microbes from other kingdoms which can have considerable effects on plant growth, disease resistance, abiotic stress tolerance and nutrient uptake (Xie et al, 2019). In CC systems, the same crop root secretions may not only affect plant growth, but also lead to simpler microbial community structure, which may negatively affect agroecosystems especially in terms of the aggravation of pathogens and soil-borne fungi. Plant-microbial interactions can have positive or negative effects on plant growth through a variety of mechanisms. Root exudates are one of the most important chemical signals in determining whether the interactions are benign or harmful (Bais et al, 2006). Identification of major differences in root exudates may be helpful in understanding changes of rhizosphere microbial community structure (Brown et al, 2020). The combination of metabolomics and microbiomics can be a powerful tool to analyze the mechanism of CC obstacles. Lin et al. (Lin et al, 2015) reported that the accumulation of allelochemicals in rhizosphere soil increased harmful microorganisms and decreased beneficial microorganisms, resulting in imbalance of microbial community structure and degradation of soil ecological function. Zhao et al. (2018) indicated that continuous coffee cultivation reduced potentially beneficial microbes in soil. Li et al. (2014b) showed that peanut root exudates promoted the growth of root rot pathogens *Fusraium oxysporum* and *Phoma* sp. and inhibited the growth of beneficial bacteria, such as *Mortierella elonate* and *Trichoderma* sp. However, the advanced metabolomics and microbiomics technologies have not been used in research of CC obstacles of alfalfa to elucidate mechanisms involved in the complex disease system.

The objectives of this research were: 1) to determine the role of continuously accumulation of autotoxic substances in CC obstacle of alfalfa; 2) to identify key metabolites causing CC obstacle and determine their effects on alfalfa seed germination and seedling growth; and 3) to evaluate the effects of key metabolites on root rot pathogens and diseases and rhizosphere microecology.

## Results

### Soil nutrients and enzyme activities

Most of the nutrient contents tested in the 7 and 14-year rhizosphere soils were lower than the 1-year rhizosphere soil, including N, P, K, B, Fe, Mn, and Mg, but Cu and Zn did not decrease with the increase of CC years (Table 1). There were significant differences in soil enzyme activities in the rhizosphere soils of different cropping years (*P* < 0.05, Fig. 1). Polyphenol oxidase, neutral phosphatase and sucrase activities decreased with the increase of CC years, while urease increased in 7-year and decreased in 14-year rhizosphere soils.

**Fig.1.**
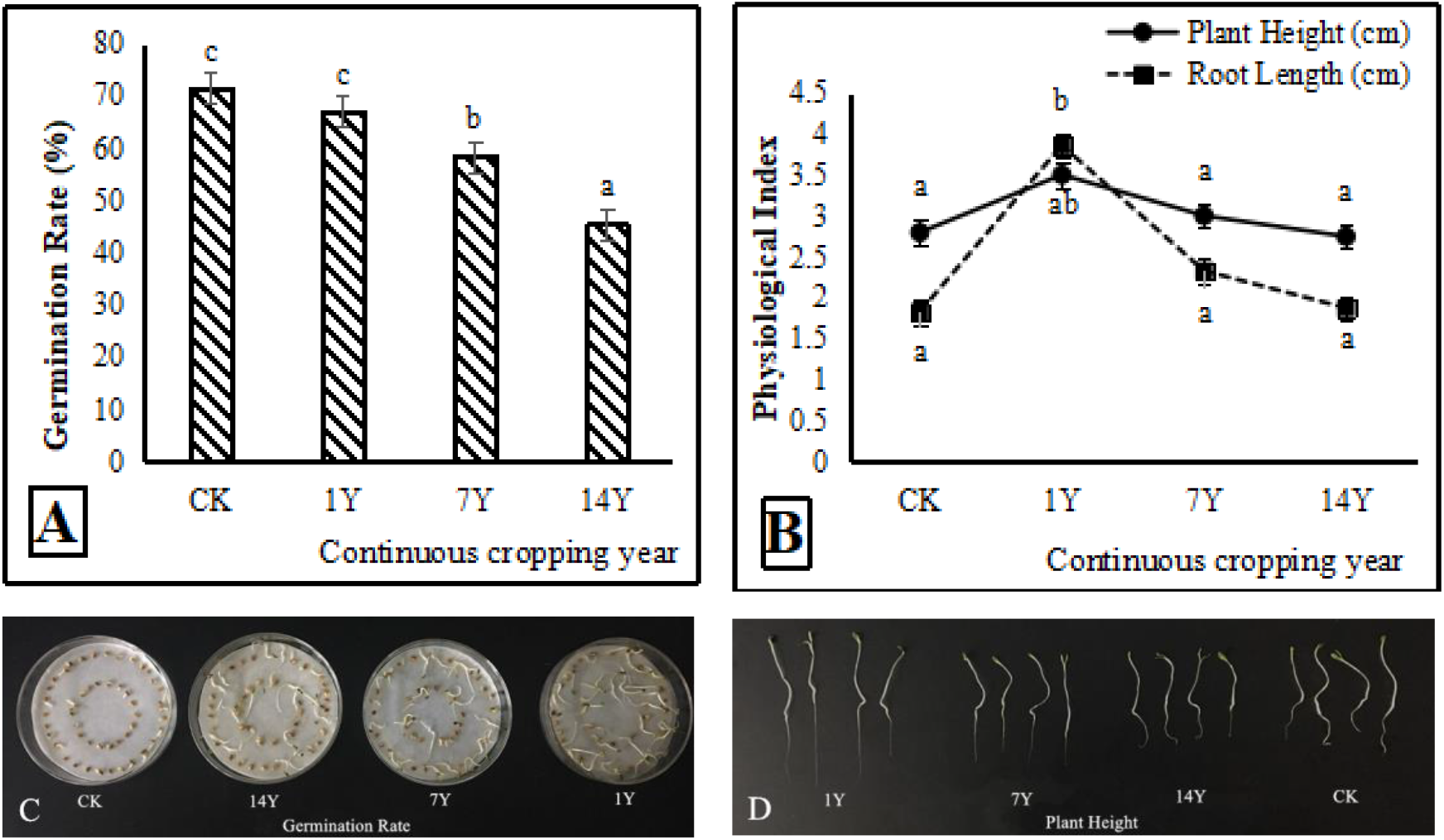
Effects of rhizosphere soil extracts from fields with different years of alfalfa continuous cropping on alfalfa seed germination (A, C) and seedling growth (B, D). Error bars indicate standard errors of the means of three repetitions. Different letters above the bars indicate significant difference according to Duncan’s multiple range test (*P* = 0.05).

**Table 1.** Nutrient content of rhizosphere soils from fields with alfalfa continuous cropping

| CC year<sup>a</sup> | Alkali-hydrolyzed N<br>(mg/kg) | Available P<br>(mg/kg) | Available K<br>(mg/kg) | Available B<br>(mg/kg) | Available Fe<br>(mg/kg) | Available Mn<br>(mg/kg) | Available Cu<br>(mg/kg) | Available Zn<br>(mg/kg) | Available Mg<br>(mg/kg) |
| --- | --- | --- | --- | --- | --- | --- | --- | --- | --- |
| 14 | 75.5 | 2.2 | 102 | 0.46 | 11.2 | 3.86 | 1 | 1.5 | 139 |
| 7 | 101 | 3.1 | 97.1 | 0.56 | 8.2 | 2.9 | 0.71 | 0.88 | 122 |
| 1 | 147.6 | 3.1 | 159 | 0.9 | 34.2 | 7.56 | 0.58 | 0.85 | 191 |

<sup>a</sup> Indicates alfalfa continuous cropping (CC) for 1, 7, and 14 years.

### Soil extracts from alfalfa rhizosphere soil on alfalfa growth

When alfalfa seeds were treated with alfalfa rhizosphere soil extracts, the germination rate in treatment with rhizosphere soil extract of alfalfa planted for 1 year was equivalent to the nontreated control, which was significantly higher than those planted for 7 and 14 years (*P* < 0.05, Fig. 2 *A* and *C*). Height of alfalfa seedlings in the 1-year treatment, but not the 7 and 14-year treatments, was significantly greater than the control (Fig. 2 *B* and *D*). However, root length of the 1, 7 and 14-year treatments was not significantly different from the control.

**Fig. 2.**
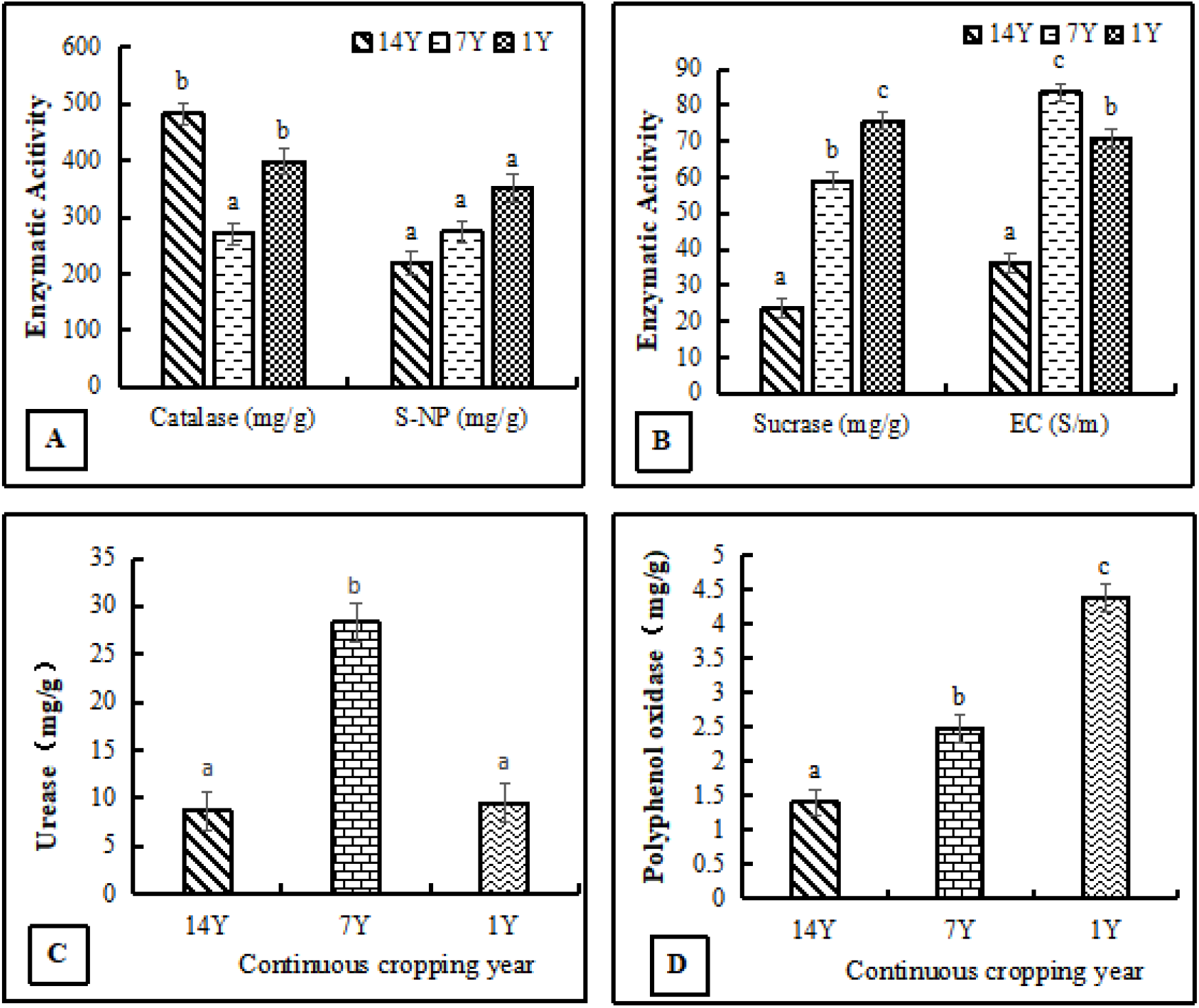
Changes of enzyme activity, electrical conduction and organic matter content in rhizosphere soils from fields with continuous cropping of alfalfa for 1, 7, and 14 years. A) Catalase and S-NP activity; B) Sucrase activity and electrical conduction; C) Urease activity; D) Polyphenol oxidase activity. Error bars indicate standard errors of the means of three repetitions. Different letters above the bars indicate significant difference according to Duncan’s multiple range test (*P* = 0.05).

### Metabolomics analysis of alfalfa rhizosphere soils

The peak with detection rate below 50% of RSD >30% was removed from the QC samples (Dunn et al, 2011). Principle Component Analysis (PCA) and least aquares-discriminant analysis (PLS-DA) were performed. A total of 161 different metabolic compounds were identified from rhizosphere soils of the 1, 7, and 14-year treatments (Fig. 3), which were classified according to their chemical property as sugar compounds (26), sugar acids (3), sugar alcohols (6), short-chain organic acids (16), long-chain organic acids (31), nuleotides (12), amino acids (14), esters (6), alcohols (16), and others (31). There were significant differences in the peak values of 58 metabolites from the rhizosphere soils. Combing VIP > 1 and independent sample T test (*P* < 0.05), there were 52 differential metabolites down-regulated and 6 differential metabolites up-regulated (Table S1). Among them, vanillic acid, p-hydroxybenzoic acid, ferulic acid, and p-courmaric acid increased significantly with the increase of CC years, and accumulation of p-hydroxybenzoic acid and p-coumaric acid was more significant with the increase of alfalfa CC years (Table 2).

**Fig 3.**
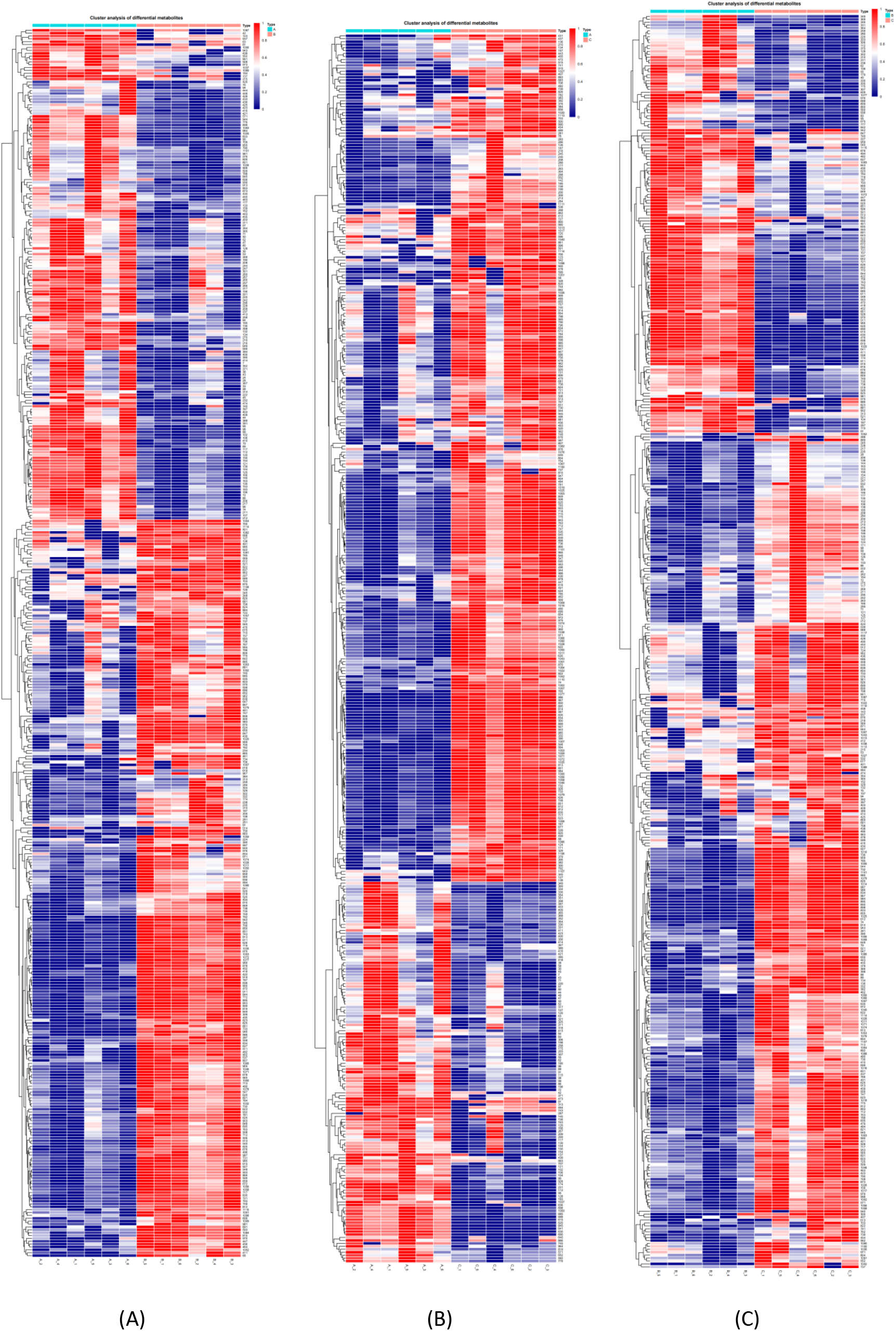
Heatmap analysis of two-year comparison of changes of alfalfa root exudates from fields with different years of continuous cropping. A) 1-year vs. 7-year; B) 7-year vs. 14-year; C) 1-year vs. 14-year.

**Table 2.** Relative content of phenolic acids from rhizosphere soils in fields with alfalfa continuous cropping

| CC year<sup>a</sup> | Vanillic acid<sup>b</sup> | p-Hydroxybenzoic acid<sup>b</sup> | Ferulic acid<sup>b</sup> | p-Coumaric acid<sup>b</sup> |
| --- | --- | --- | --- | --- |
| 1 | 0.050 ± 0.008 a | 0.127 ± 0.014 a | 0.031 ± 0.004 a | 0.056 ± 0.011 a |
| 7 | 0.068 ± 0.007 b | 0.171 ± 0.013 b | 0.026 ± 0.004 a | 0.079 ± 0.009 b |
| 14 | 0.070 ± 0.006 b | 0.212 ± 0.017 c | 0.046 ± 0.012 b | 0.097 ± 0.011 c |

<sup>a</sup> Indicates alfalfa continuous cropping (CC) for 1, 7, and 14 years.
<sup>b</sup> Different letters in the column indicate significant difference according to Duncan's multiple range test ( $P = 0.05$ ).

The effects of four phenolic acids on alfalfa plant were evaluated at concentrations of 10, 25, 50 and 100 μg/mL. Inhibitory effect on seed germination and plant growth was as follows: p-coumaric acid > ferulic acid > vanillic acid and p-hydroxybenzoic acid, with higher concentrations having greater inhibition than a lower concentration (Fig. 4). P-coumaric acid reduced root length significantly at all concentrations, compared to the control, and the other three compounds reduced root length significantly at higher concentrations (Fig. 4 *E-H*). Higher concentrations of ferulic acid and p-hydroxybenzoic acid reduced plant height significantly compared to the control, but vanillic acid did not reduce plant height at all concentrations (Fig. 4*E*).

**Fig. 4.**
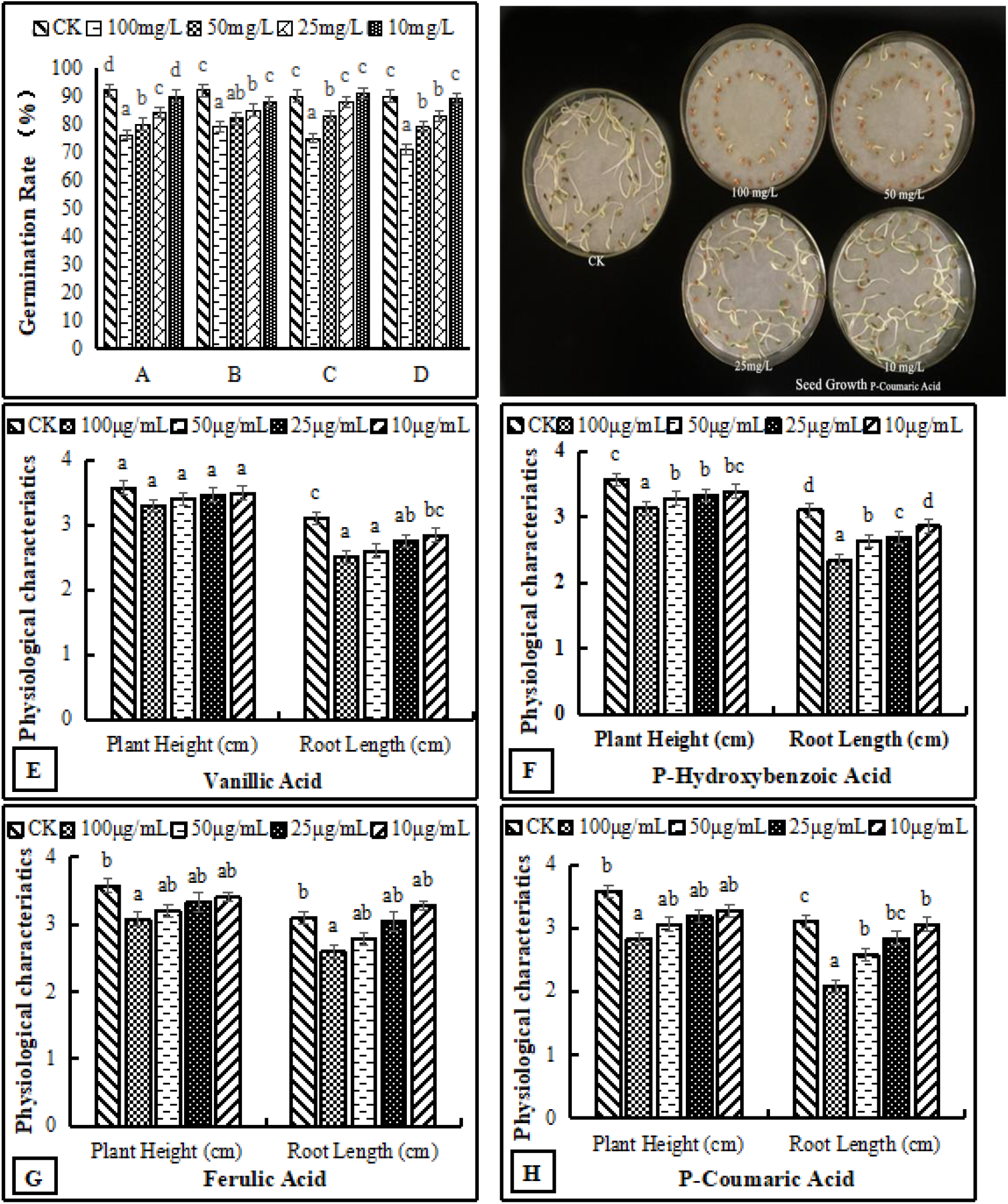
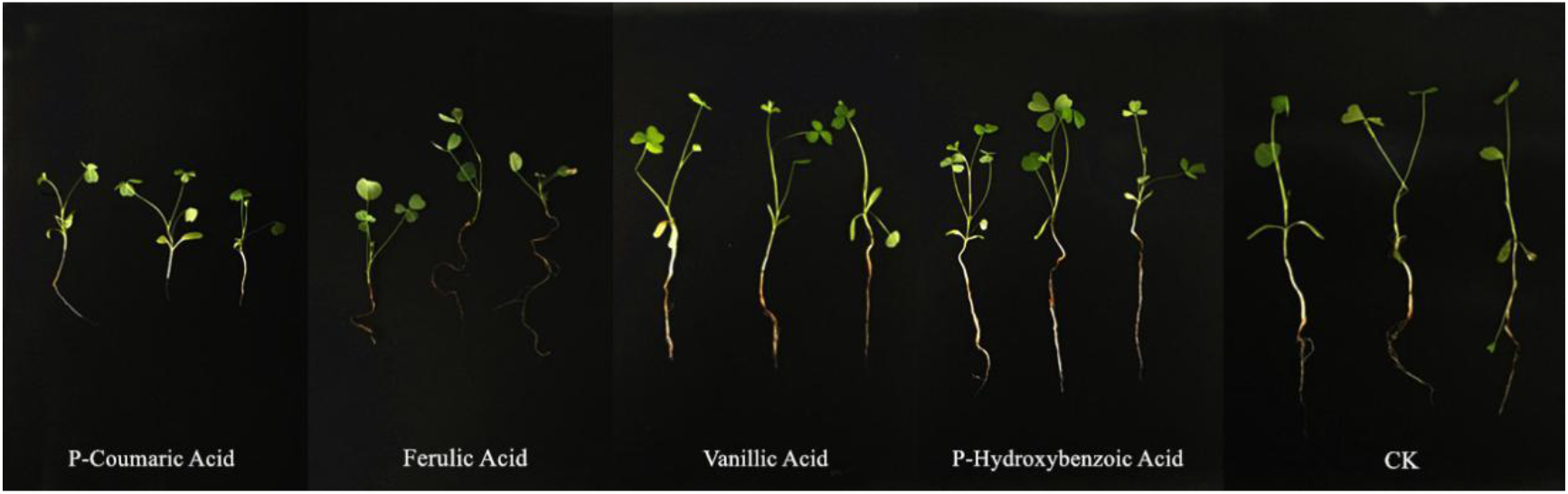
Effect of different concentrations of phenolic acids on alfalfa germination rate, root length and plant height (E, F, G, H). A) Vanillic Acid; B) P-Hydroxybenzoic Acid; C) Ferulic Acid; D) P-Coumaric Acid. Error bars indicate standard errors of the means of three repetitions. Different letters above the bars indicate significant difference according to Duncan’s multiple range test (*P* = 0.05).

### Key metabolites from alfalfa rhizosphere soils on alfalfa root rot pathogens

Three fungal pathogens *Fusarium trinctum*, *F. acuminatum* and *F. oxysporum* that cause alfalfa root rot were used to assess the relationship between metabolites of alfalfa rhizosphere and root rot of alfalfa. Extracts of rhizosphere soil in 14-year alfalfa CC significantly enhanced mycelial growth of *F. trinctum*, *F. acuminatum* and *F. oxysporum* when used at 1%, 5% and 10% compared to 1-year and the water control (Fig. 5). Extracts of rhizosphere soil in 7-year alfalfa CC also enhanced mycelial growth of the pathogens significantly when used at higher concentrations. The effects of four phenolic acids on mycelial growth of the fungal pathogens were evaluated. They enhanced mycelial growth with effects in the following order: p-coumaric acid > ferulic acid > vanillic acid >p-hydroxybenzoic acid (Fig. 6). Higher concentrations of the phenolic acids had greater effects in enhancing mycelial growth of the fungal pathogens. When tested at 10, 25, 50 and 100 µg/mL, the four phenolic acids at higher concentrations also significantly enhanced conidial germination and production of *F. acuminatum* and *F. oxysporum* (Fig. 7). Among the phenolic acids, p-coumaric acid had the greatest promoting effect on conidial germination and production. When inoculated with *F. trinctum*, *F. acuminatum* and *F. oxysporum* in greenhouse studies, severity of alfalfa root rot treated by p-coumaric acid increased significantly with 50 μg/mL having greater effect than 10 μg/mL (Fig. 8). Growth and development of seedlings were also suppressed by p-coumaric acid.

**Fig. 5.**
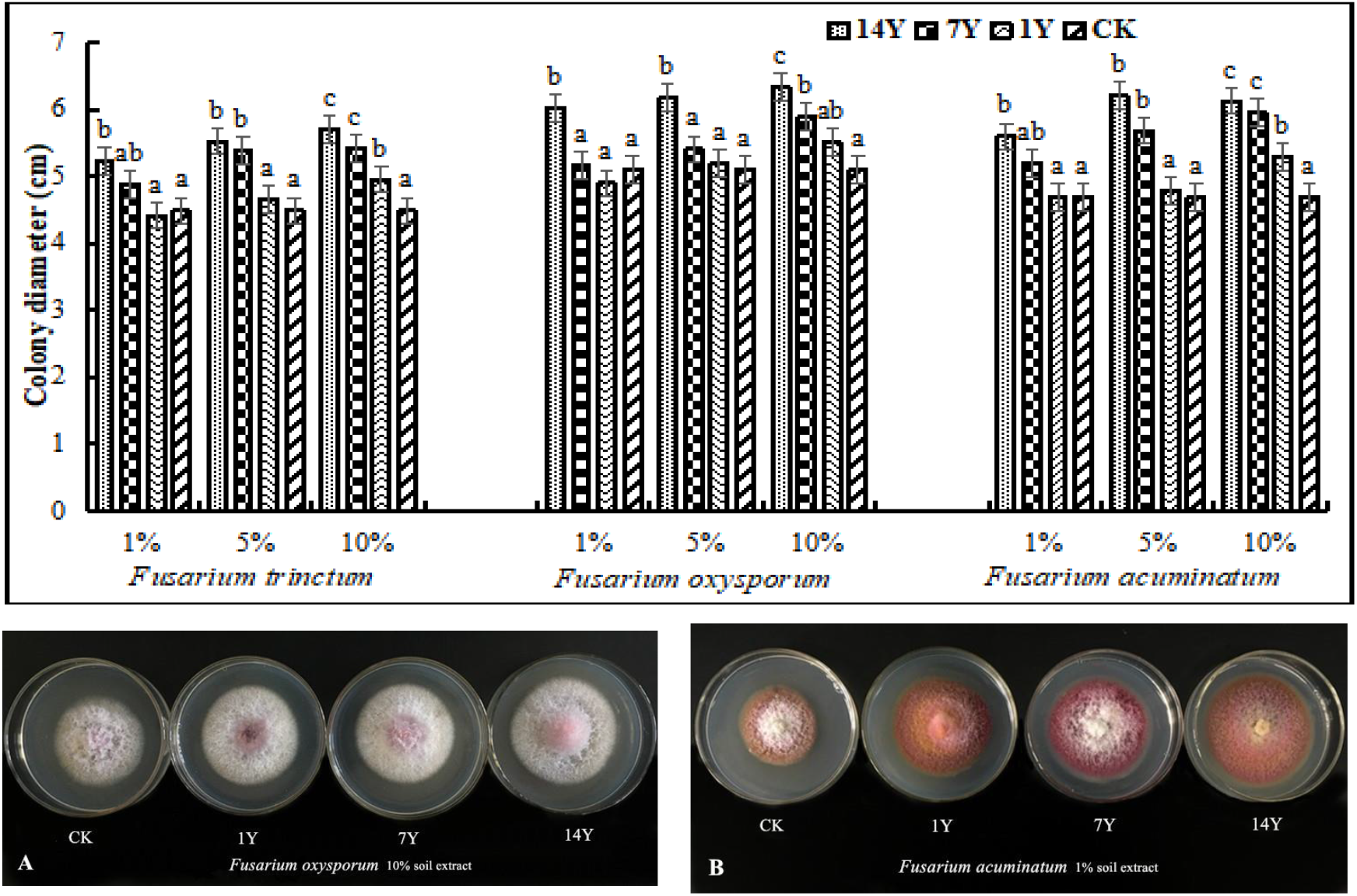
Effect of rhizosphere soil extracts (final concentration V/V: 1%, 5% and 10%) from fields with alfalfa continuous cropping for 1, 7, and 14 years on mycelial growth of *Fuasrium oxysporum*, *F. trinctum* and *F. acuminatum* that cause alfalfa root rot. A) Effect of 10% soil extract on mycelial growth of *F. oxysporum*; B) Effect of 1% soil extract on mycelial growth of *F. acuminatum.* Error bars indicate standard errors of the means of three repetitions. Different letters above the bars indicate significant difference according to Duncan’s multiple range test (*P* = 0.05).

**Fig. 6.**
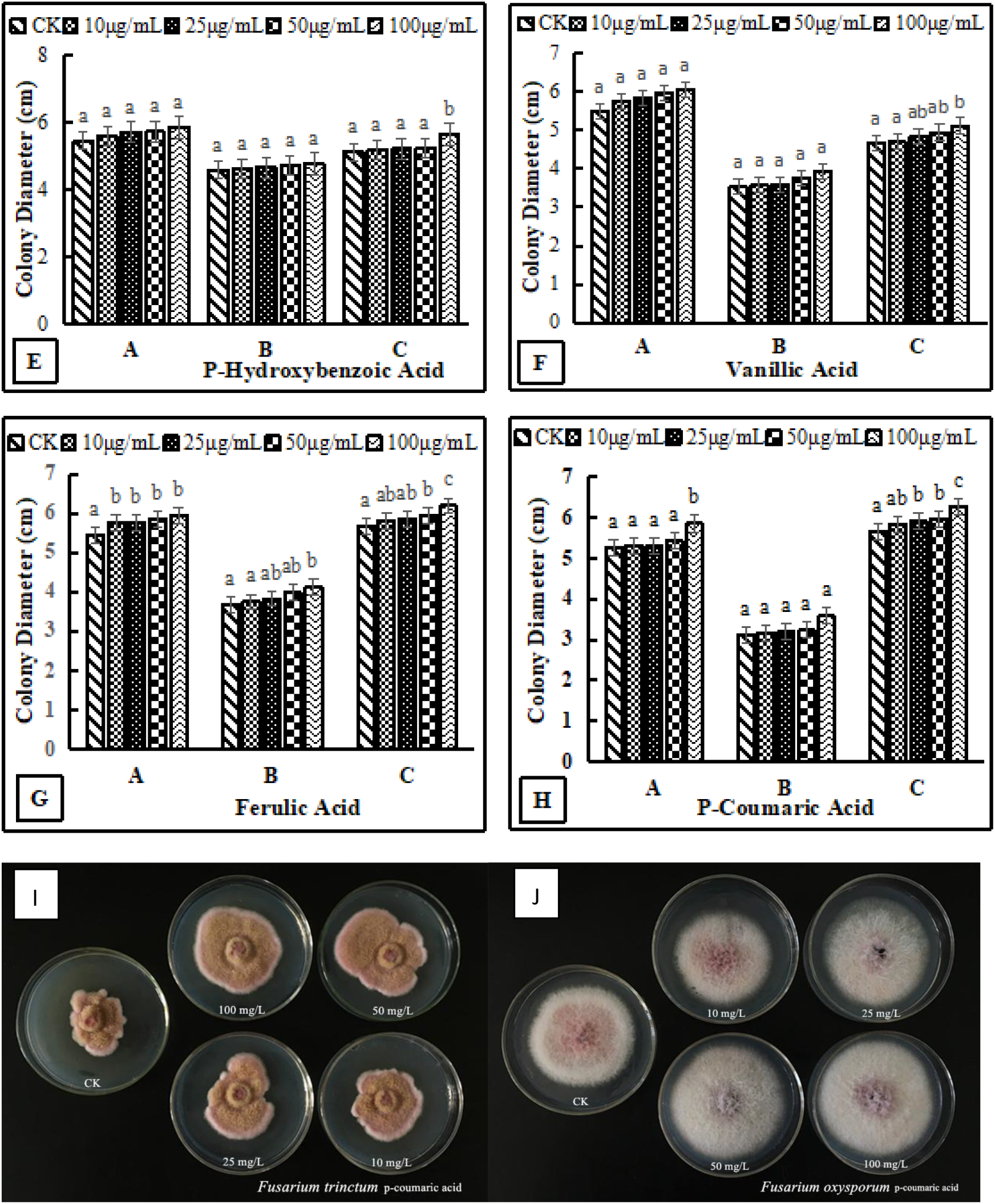
Effects of different concentrations of phenolic acids on mycelial growth of fungal pathogens that cause alfalfa root rot. A, B and C indicate *Fuasrium oxysporum*, *F. trinctum* and *F. acuminatum*, respectively (E, F, G, H). Error bars indicate standard errors of the means of three repetitions. Different letters above the bars indicate significant difference according to Duncan’s multiple range test (*P* = 0.05). I) Effect of p-coumaric acid on mycelial growth of *F. trinctum*; J) Effect of p-coumaric acid on mycelial growth of *F. oxysporum*.

**Fig. 7.**
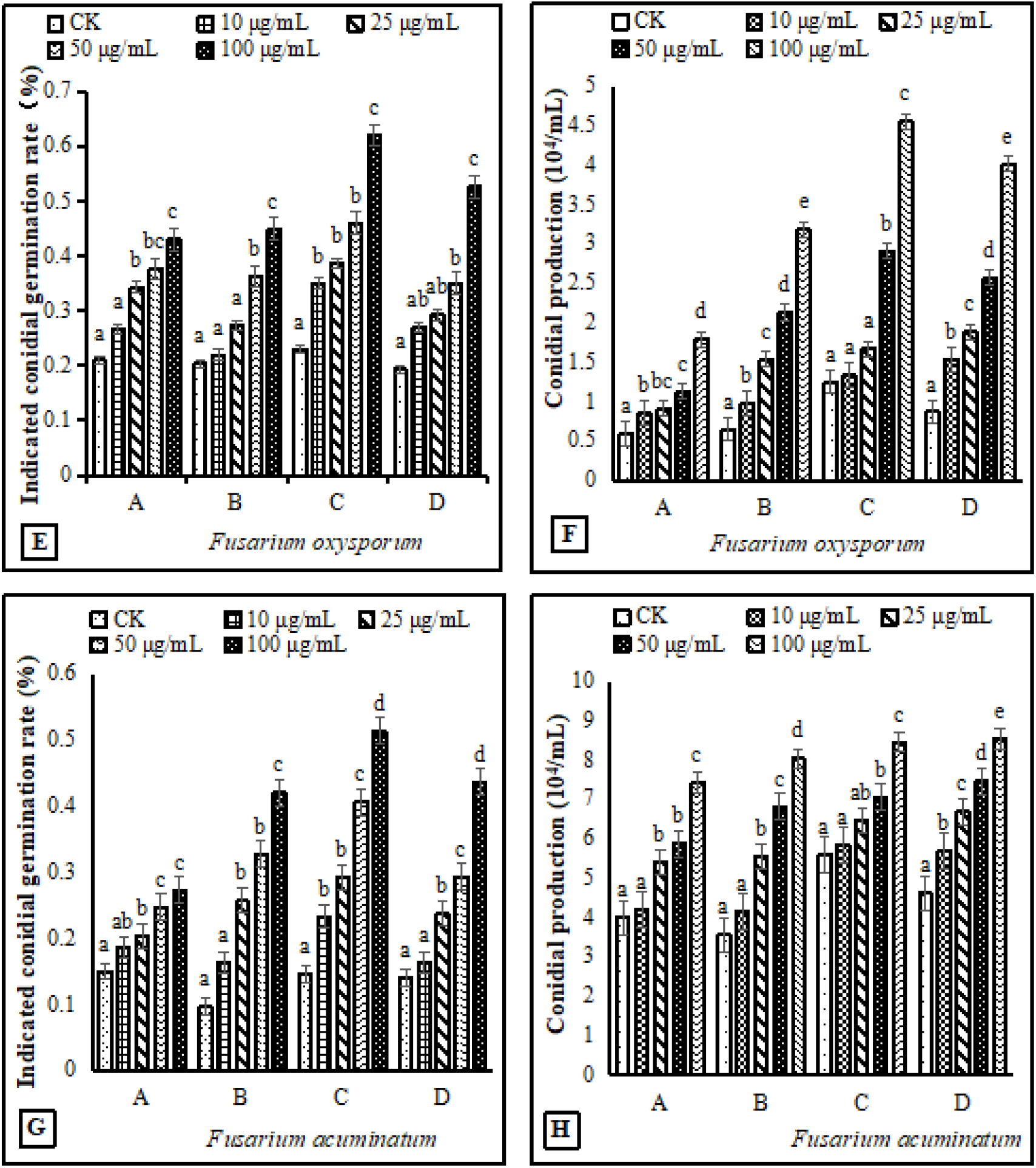
Effects of phenolic acids on conidial germination and production of *Fusarium oxysporum* (E, F) and *F. acuminatum* (G, H) that cause alfalfa root rot. A, B, C and D indicate p-hydroxybenzoic acid, vanillic acid, ferulic acid and p-coumaric acid, respectively. Different letters above the bars indicate significant difference according to Duncan’s multiple range test (*P* = 0.05).

**Fig. 8.**
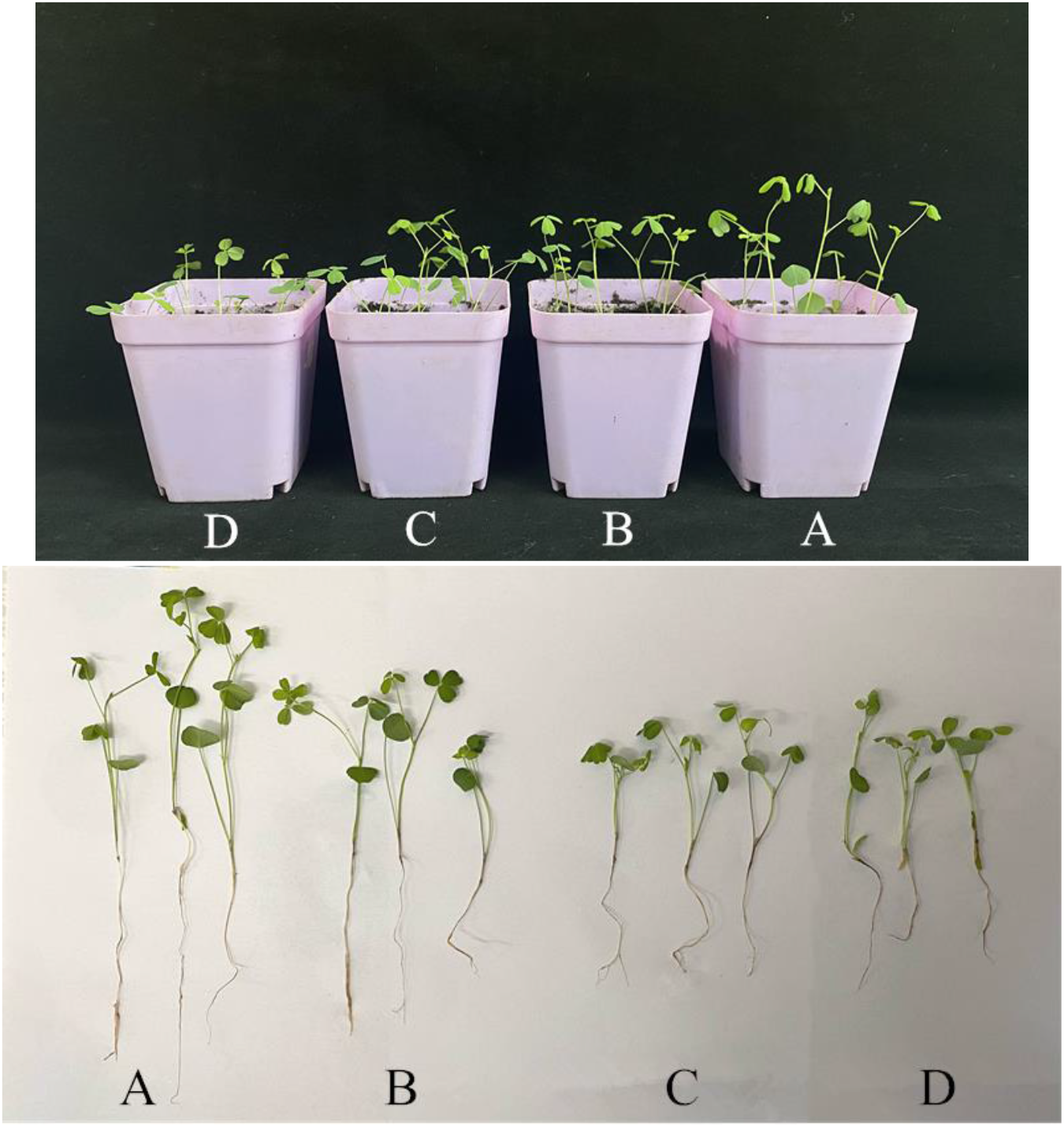
Effect of p-coumaric acid on alfalfa seedling and root rot. A) Inoculated with water as control; B) Inoculated with mixed spore suspensions of *Fusarium trinctum*, *F. acuminatum* and *F. oxysporum*; C) Inoculated with mixed spore suspensions of the three *Fusarium* spp. and 10 μg/mL p-coumaric acid; D Inoculated with mixed spore suspensions of the three *Fusarium* spp. and 50 μg/mL p-coumaric acid.

### Alfalfa CC on alfalfa rhizosphere microecology

Microbial community sequences of the rhizosphere soil samples were analyzed on Illumina Miseq PE300 platform and divided by OTUs. Principle Component Analysis (PCoA) based on detected OTUs showed that there were significant differences in community composition of the rhizosphere soils, and Anova analysis showed that there were significant differences among groups (*P* < 0.05, Fig. 9 *D1* and *E1*). The diversity of bacterial community was high and the number of OTUs decreased with the CC increase (Fig. 9 *D2*). The diversity of bacterial community of 14-year CC was the lowest, while the diversity of fungal community of 1-year cropping was the lowest (Fig. 9 *E2*). Small changes in root exudates can lead to changes in soil microbial structure (Ling et al, 2011). This change could lead to soil microecological damage as well as disease and pest problems (Liu et al, 2020). These results seem to indicate that CC can change the microbial composition of soil from bacterial-dominated to fungal-dominated (Lin et al, 2015).

**Fig. 9.**
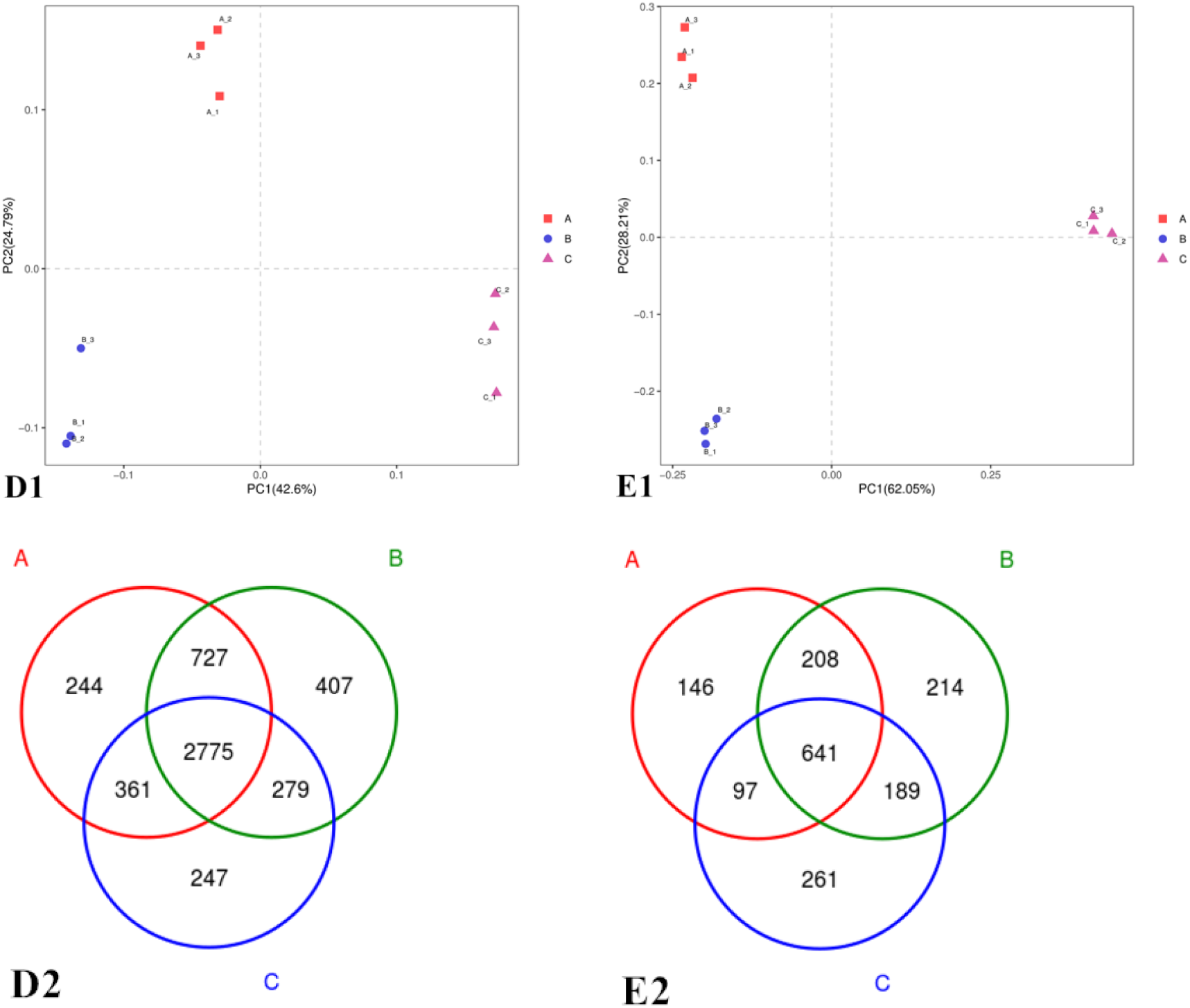
Principal coordinate analysis (PCoA) showing the similarity of rhizosphere microbial community in fields with continuous cropping of alfalfa for 1 year (A), 7 years (B), and 14 years (C). D1) bacteria, and E1) fungi. D2) Venn diagram of bacteria, and E2) Venn diagram of fungi.

Most OTUs of bacteria were *Actinobacteria* (35.55%, 37.54%, and 40.28% for 1-year, 7-year and 14 year CC respectively), *Acidobacteria* (24.07%, 18.49%, and 19.53% for 1-year, 7-year and 14-year) and *Proteobacteria*, and a small proportion belonged to *Gemmatimonadetes*, *Verrucinobacter*, *Nitrospiraceae*, and *Bacteroidetes* (Fig. 10*D*). *Bacillus* is considered an important soil conditioner and can be used as a biological control agent (Li et al,, 2018). In our present study, the abundance of *Bacillus* decreased in years and then increased in 14 years. Most OTUs of fungi belonged to *Ascomycota* (56.14%, 65.57%, and 73.72% for 1, 7, and 14-year CC, respectively), *Basidiomycota*, *Mortierellomycota*, *Glomeromycota*, and *Chytridiomycota* (Fig. 10*E*).

**Fig. 10.**
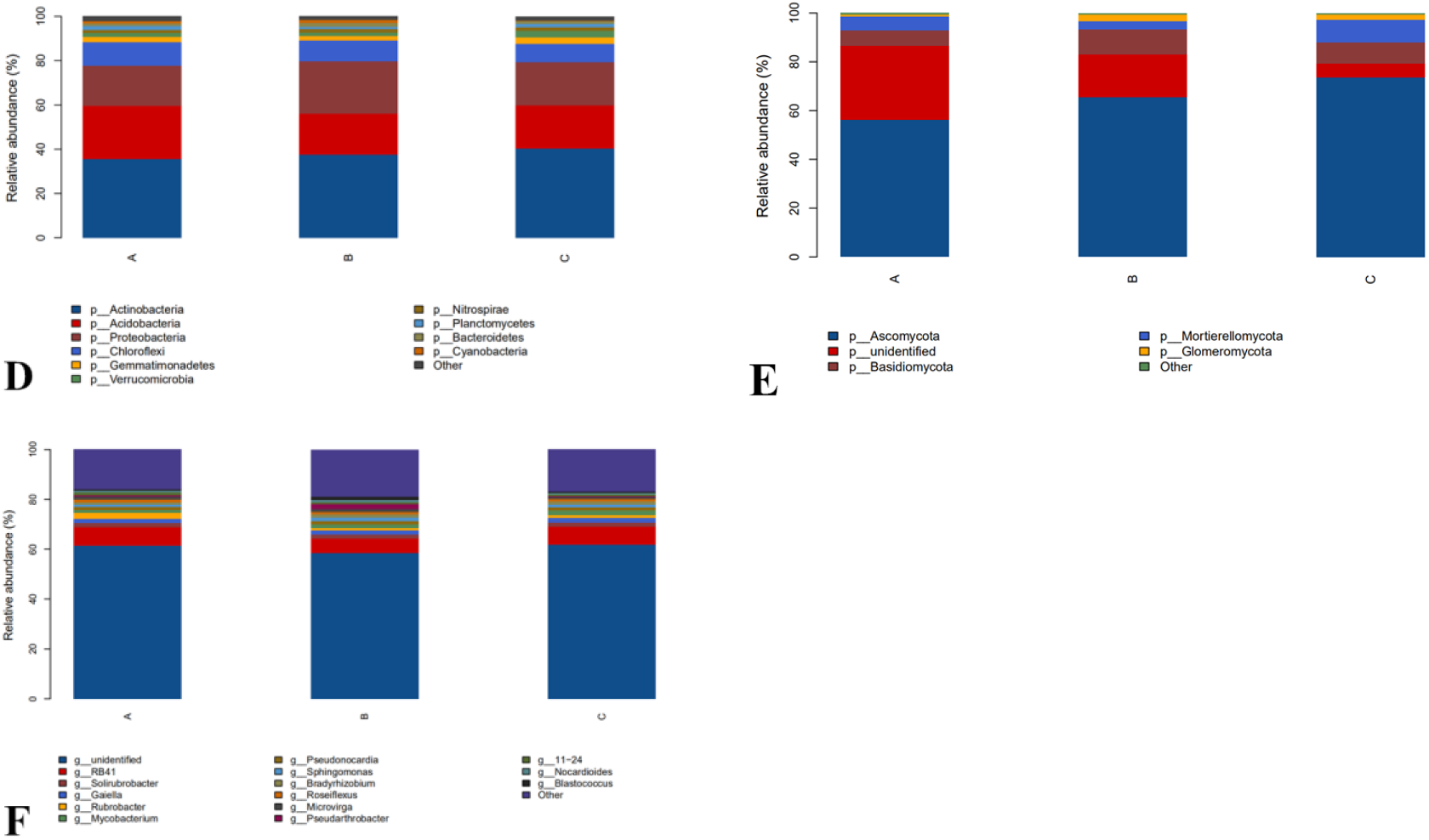
The relative abundance (%) of major bacteria (D) and fungi (E) phylum, and bacterial genus (F), in the microbial community of alfalfa rhizosphere soils from fields with continuous cropping of alfalfa for 1, 7, and 14 years.

### P- coumaric acid on rhizosphere microbial communities

In order to verify the correlation between microbial population changes and metabolites secreted in the rhizosphere, alfalfa rhizosphere soil was treated with p-coumaric acid, the most influential metabolite secreted by alfalfa. The results showed that significant changes occurred in the bacterial and fungal communities (Fig.11 *D1* and *E1*). The number of OTU in bacterial and fungal communities decreased when treated with 10 μg/mL p-coumaric acid but increased when treated with 50 μg/mL p-coumaric acid. The number of OTU in soil bacterial community treated with p-coumaric acid was less than that of the control (Fig.11 *D2* and *E2*). The biodiversity of bacterial community was the highest when treated with 10 μg/mL of p-coumaric acid, the lowest with 50 μg/mL, and the diversity of soil fungal community increased gradually with the increase of p-coumaric acid concentration, which was consistent with the previous results of continuous cropping. However, the number of microorganisms in alfalfa rhizosphere soil in the pots treated with p-coumaric acid was lower than that in the field.

**Fig. 11.**
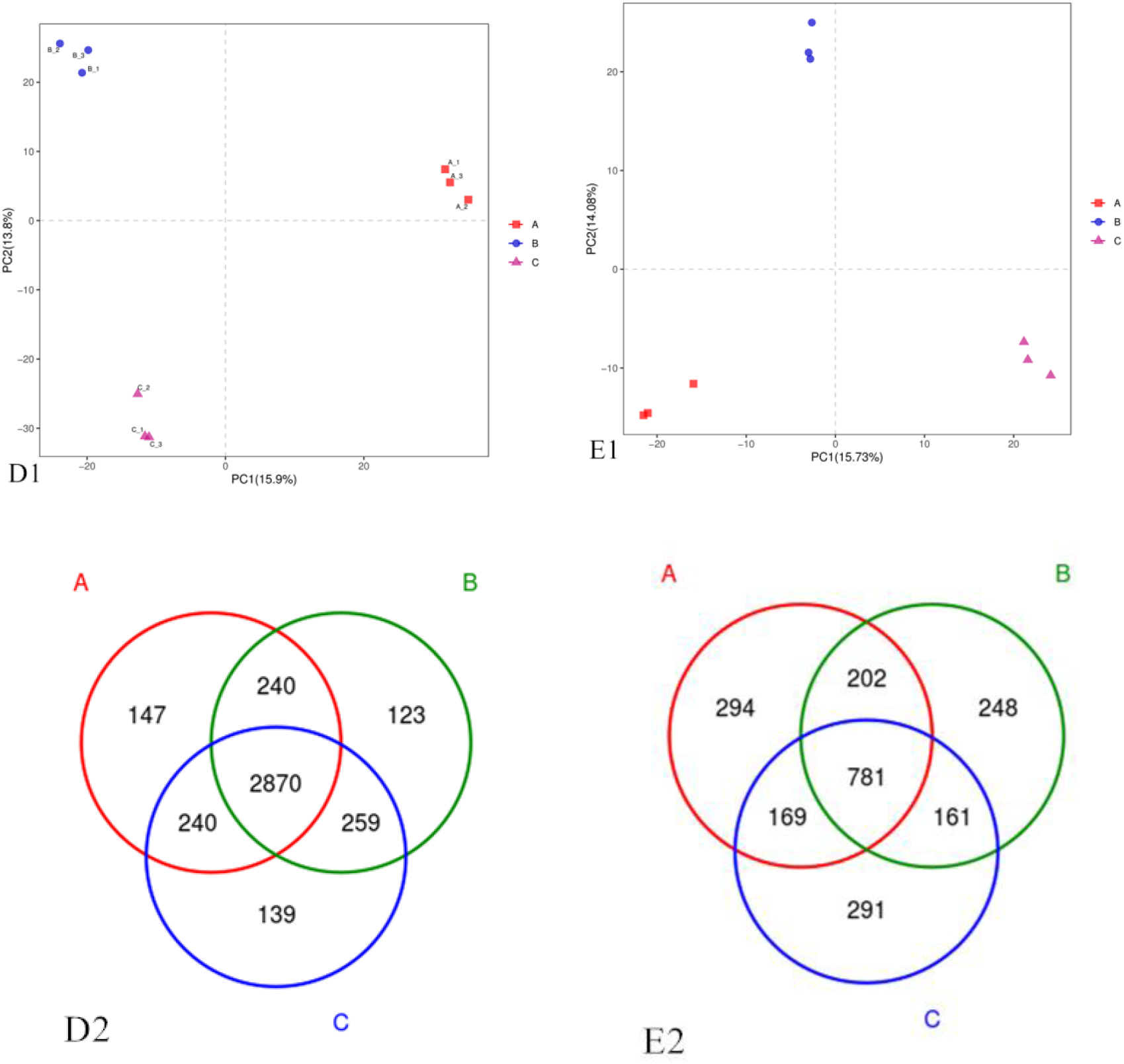
Principal coordinate analysis (PCoA) showing the similarity of rhizosphere microbial community in fields with A (nontreated), B (treatment with 10 μg/mL p-coumaric acid), C (treatment with 50 μg/mL p-coumaric acid). D1) bacteria, and E1) fungi. D2) Venn diagram of bacteria, and E2) Venn diagram of fungi.

Analysis of microbial species and functions indicated that the *Gemmatimonadetes* decreased with increasing concentration of p-coumaric acid (Fig.12*D*). The relative abundance of bacterial species decreased to different degrees, with 8 main bacterial species in the top 20 species showing a decreasing trend, 6 bacterial groups showing the trend of decreasing when treated with 10 μg/mL p-coumaric acid and increasing with 50 μg/mL, and 5 groups of bacteria showing the trend of increasing with 10 μg/mL and decreasing with 50 μg/mL (Fig. 12*F*). *Sphingomonas* increased with the addition of p-coumaric acid, which was consistent with the experimental results described above. *Sphingomonas* and *Gemmatimonas* were related to nitrogen metabolism and transformation, as well as the change of NH_4_^+^-N and N_2_O contents (Nathanae et al, 2009; Shen et al,2014, Li et al, 2017). *Candidatus Solibacter* is able to decompose organic matter, and the results showed it decreased with the increase of p-coumaric acid concentration (Zak et al, 1996). *Bryobacter* promoted soil carbon cycle (Li et al, 2012), which showed a trend of increasing at first and then decreasing, and was lower in the p-coumaric acid treatment (50 μg/mL) than the control soil (Fig.12*E*).

**Fig. 12.**
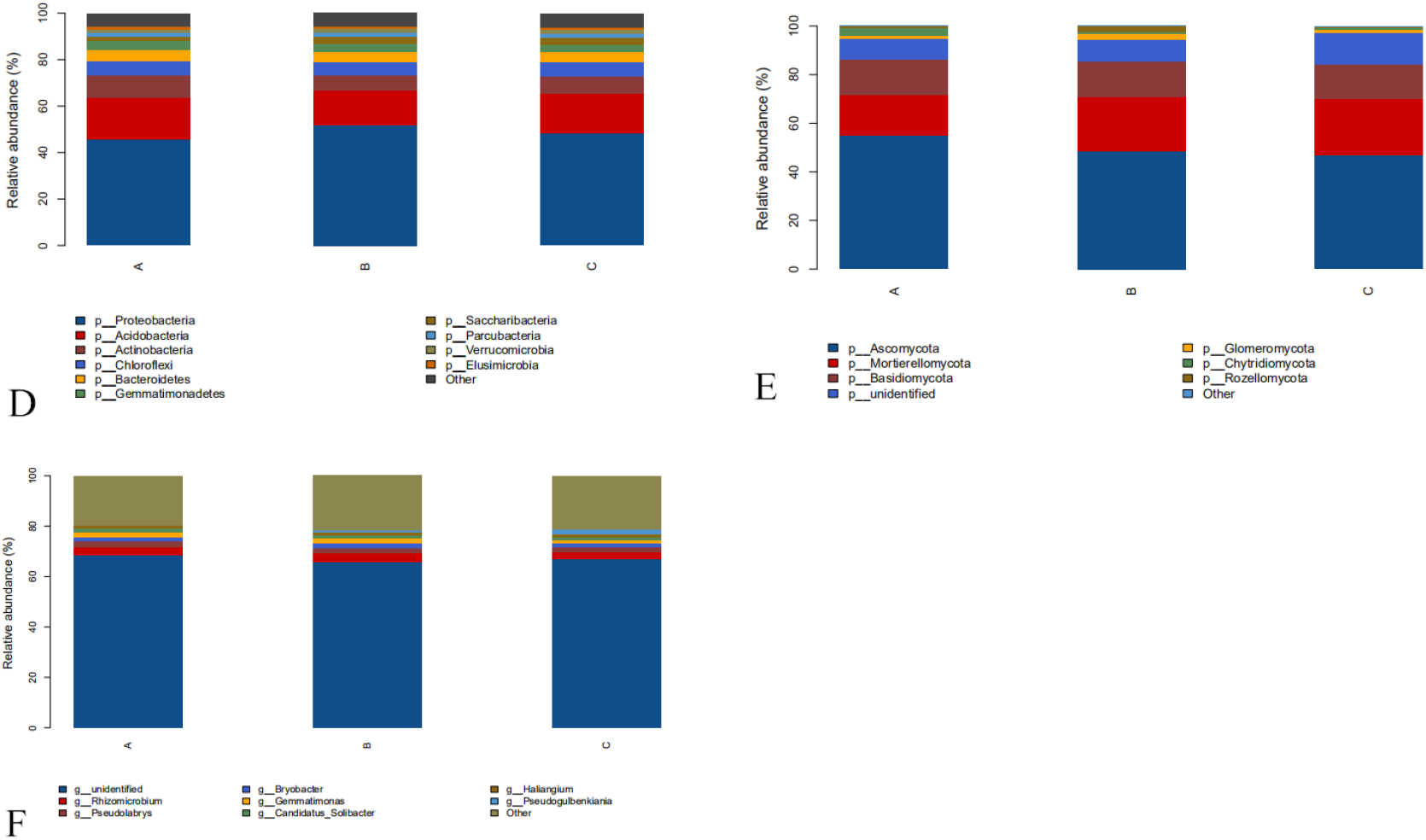
The relative abundance (%) of major bacteria (D) and fungi (E) phylum, and bacterial genus (F), in the microbial community of alfalfa rhizosphere soils from fields with A) nontreated, B) treatment with 10 μg/mL p-coumaric acid, and C) treatment with 50 μg/mL p-coumaric acid.

## Discussion

The soil enzyme is a key criterion for evaluating different residue covering approaches, as it is an important indicator of soil quality and function. Sucrase can reflect the conversion ability of organic carbon (Cantarella et al, 2018). As the CC years increase, the conversion of organic carbon gradually decreases, which is consistent with studies of Liu et al. (2017). Soil urease reflects the transformation ability of soil organic nitrogen to available nitrogen and the supply ability of soil inorganic nitrogen (Tawaraya et al, 2014). Phosphatase can catalyze the hydrolysis of soil monophosphate to form inorganic phosphorus, which can be absorbed by plants, and soil phosphatase activity can be used to characterize the state of soil phosphate (Hu et al, 2015). Our present study indicated that the activity of soil phosphatase transformation decreased with the increase of CC years. In the meantime, the content of phosphorus in soil decreased with the increase of CC years. Soil catalase activity indicates its ability to remove the toxicity of hydrogen peroxide, which could reflect soil quality and the total metabolic activity of soil microorganisms (Zhang et al, 2012). Overall, soil quality became worse with the increase of CC years of alfalfa.

Development of alfalfa seedlings needs the support of external nutrition. In the study plant height and root length from seeds treated with rhizosphere soil extracts decreased with the increase of CC years, which might be because soil nutrient condition gradually became worse. However, theoretically nutrition for seed germination is provided by the nutrition stored by the seed itself and there is little need for extra nutrition. The richness of soil nutrients only affects the growth of seedlings after germination but does not affect seed germination rate (Meng et al, 2006). Our present study provides two evidences that do not support the view that the main factors of alfalfa CC obstacle were the lack of soil nutrients. One was that seed germination does not need external nutrients, and at least the germination rate of alfalfa seeds treated by soil extracts from the 1, 7, and 14-year CC should be similar to that of the water treatment control. However, it is interesting that seed germination rate decreased significantly with the increase of CC years. Another evidence was that nutrients in the soil extracts from the three alfalfa cropping years were richer than the water control. In theory the soil extracts of the three cropping years should have significant effects on the growth of alfalfa seedlings compared to the control. However, our results showed that the effects of soil extracts from CC for 7 and 14 years on seedling growth were similar to that of the control. These results indicated that lack of nutrients in alfalfa rhizosphere soil was not the key factor resulting in alfalfa CC obstacles.

As our results, the accumulation of phenolic acids in the soil after CC significantly affects the rhizosphere ecosystem, for instance, by inducing changes in microbial populations, soil enzyme activity and nutrient cycling (Halvorson et al, 2009; Chen et al, 2020). Many perennial and annual crop species are threatened by CC problems associated with reduced plant growth and vigor as well as reduced crop yields and quality (Chen et al, 2012; Li et al, 2018; Li & Liu, 2018). In fact, previous studies have shown that phenolic acids are a major secondary metabolite of CC disorder (Muscolo & Sidari, 2006). In studies on cucumber, strawberry, tobacco and Rehmannia, soils continuously planted were found to contain self-toxic substances, phenolic acid, which repressed growth of the same plant (Li et al, 2012; Wu et al, 2009; Chen et al, 2011; Li et al, 2015b). With the extension of CC years, the concentration of phenolic acid in the soil increased gradually (Huang et al, 2013). Qu and Wang (Qu & Wang, 2008) reported that two phenolic acids from soybean root exudates, 2,4-di-tert-butylphenol and vanillic acid, had significant negative effect on microbial communities and soybean monoculture. Li et al. (2012) also found that the extracts from rhizosphere soil of Ginseng CC had significant inhibition allelopathy on the growth of radicle. These are consistent with our findings about the inhibitory effects of the phenolic acids on alfalfa. With the increase of alfalfa CC years, the contents of four phenolic acids (vanillic acid, p-hydroxybenzoic acid, ferulic acid and p-courmaric acid) increased significantly (Table 2). Therefore, we thought that the occurrence of alfalfa CC obstacle may be related to the phenolic acids.

Crop roots respond to pathogen infections by changing the amount and composition of root exudates (Lanoue et al, 2010). Studies showed that cucumber CC usually led to the accumulation of soil-borne pathogens such as *Fusarium* spp. (Zhou & Wu, 2012). In alfalfa CC, we also found that root rot caused by diverse pathogens increased with the increase of alfalfa CC years. Wu et al. (2015) reported that a mixture of phenolic acids promoted the growth of *F. oxysporum* hyphae, spore formation and production. Zhou et al. (2012) indicated that the amount of p-coumaric acid from cucumber could increase the number and population density of *F. oxysporum* in soil and disease incidence in the field. A number of studies also demonstrated that CC significantly increased levels of fungal pathogens causing root diseases of *Rehmannia glutinosa*, soybean and cucumber (Wu et al, 2015; Guo et al, 2011; Zhou & Wu, 2012a). These studies are in agreement with our research that alfalfa root rot was getting more severe with the increase of CC years. Our present study further confirmed that metabolites secreted by alfalfa rhizosphere, such as p-coumaric acid, had strong effects on alfalfa root rot and may be the key factor of alfalfa CC obstacle.

Soil bacteria are responsible for decomposing organic matter into inorganic matter and thus maintaining soil fertility, so we did a differential analysis of soil bacterial flora for different years of CC (Fig. 10*F*). The results show that *Gaiella*, *Pseudonocarida*, *Mycobacterium* and *Bradyrhizobium* in the soil samples increased gradually with the increase of CC years, and *Solirubrobacter* decreased with the increase of CC years. The relative abundance of *Sphingomonas* was less than 1% in 1-year and more than 1.4% in 14-year treatments, showing an increase with CC. Other studies have shown that *Sphingomonas* was a nutrient-poor nanobacterium, which could adapt to heterotrophic growth under conditions of nutrient depletion (Williams et al, 2009; Kämpfer et al, 2002; Pooja et al, 2010). In this paper, the increase of *Sphingomonas* indicates that after 14 years of continuous cultivation, the rhizosphere soil presented a state of nutrient depletion. This is consistent with the results of Chen et al. (2020). The relative abundance of bacteria RB41 was higher than 7.2% in 14 years of CC, which was higher than in 1 and 7-years. The acid bacteria may play an important role in remaining the metabolism of soil under long-term low nutrient stress, and could degrade the polymer of plant residues (Aislabie et al, 2006; Fan et al, 2018). *Bradyrhizobium* is a diazotrophic bacterium in soil, which plays an important role in nodule formation, ammonia production and nitrogen fixation symbiosis in legume roots (Masuda et al, 2016; Shiro et al, 2016; Saeki et al, 2017; Siqueira et al, 2017). The difference of relative abundance of *Bradyrhizobium* in three different CC years was significant, which was higher in 14-year CC than in 1-year. Two fungi *Gibberella* and *Metarhizium* were 11.5% in soil after one year of CC and 1-2% after 14 years of CC. In the CC process, the bacterial community in soil decreased significantly.

It appeared that the change of alfalfa rhizosphere microbial communities treated with p-coumaric acid was basically consistent with that of alfalfa CC obstacle. There were some differences between results in the potted alfalfa study and the actual field conditions, which was probably due to other secondary autotoxic substances. Overall, alfalfa CC had an impact on soil microbial communities, and the accumulation of autotoxins in rhizosphere soils increased harmful microorganisms and decreased beneficial microorganisms (such as *Gemmatimonadetes* and *Sphingomonas et al*), resulting in imbalance of microbial community and degradation of soil ecological function.

And the results of the four phenolic acid treated seeds and pathogens were similar to those treated by soil extracts. The occurrence of CC obstacle was directly related to the four phenolic acids secreted by alfalfa rhizosphere. Among them, the effects of ferulic acid and p-coumaric acid on alfalfa seed germination and mycelium growth were significant. The effects of p-coumaric acid on alfalfa seedling development were obvious, and the effects of ferulic acid on spore production and germination of pathogenic fungi causing alfalfa root rot were significant. Bi et al. (2010) reported that nine phenolic acids, such as p-hydroxybenzoic acid, coumaric acid, and ferulic acid, were detected in commercially grown Ginseng rhizosphere soils, which could inhibit the growth of radicles and buds. Zhou &Wu (2012a) found p-coumaric acid, an autotoxin of cucumber, increased *F. oxysporum* f. sp. *cucumerinum* population densities in soil and the severity of Fusarium wilt under field conditions. Wu et al. (2015) indicated that phenolic acid mixtures promoted hyphal growth, spore formation and production of *F. oxysporum* that causes wilt disease of *Rehmannia glutinosa*. Tao et al. (2018) reported that both p-hydroxybenzoic acid and ferulic acid could inhibit alfalfa seed germination and seedling development. Therefore, we think that the four phenolic acids secreted in the rhizosphere of alfalfa were the main factors causing severe alfalfa root rot in the alfalfa CC obstacle.

In this study, we found that the main factors causing CC obstacle were not the lack of nutrients or water in alfalfa rhizosphere soil. Based on metabolomics and microbiology analysis, the effects of certain key metabolites, including p-coumaric acid, ferulic acid and other phenolic acids, on alfalfa seed and seedling growth and root rot pathogens were basically consistent with the influence of CC obstacles in the field. In addition, with the increase of CC years, the microbial community in soil changed from fungal to bacterial, and beneficial microorganisms decreased with the increase of CC years. The effects of the key metabolites from alfalfa rhizosphere on alfalfa seed germination, seedling growth and root rot were further verified, which resulted in similar alfalfa performance as in CC obstacles. Among these key metabolites, the autotoxicity of p-coumaric acid was the strongest. This study fully proved that the continuous accumulation of autotoxic substances in the rhizosphere was the key factor of alfalfa CC obstacle.

## Materials and Methods

### Soil Sampling

The study was conducted on an experimental farm located in Dörbets, Daqing, Heilongjiang, Northeast China (124.25′ E, 46.30′ E) with a continental monsoon climate. The soil type was sandy loam and average depth of topsoil was about 30 cm. No location had history of hardpan. Soil samples were collected in October 2019 using the five-spot-sampling method from the rhizosphere of alfalfa plants grown in the fields with a history of alfalfa CC for 1, 7 and 14 years. The soil samples, thoroughly homogenized through a 20-mesh sieve to remove root debris, were placed in sterile bags and then transferred to liquid nitrogen and stored in ice boxes. The samples were transported to the laboratory and stored at -80℃ for metabonomic and microbiological analysis. In the meantime, a portion of the soil samples was dried for analysis of soil properties. Each treatment had three replicate samples.

### Assessment of soil properties

Physic-chemical properties of the rhizosphere soils, including available P (AP), available N (N), Fe, Mn, Cu, Zn and EC values, were analyzed at the Soil and Fertilizer Testing Center of Heilongjiang Academy of Agricultural Sciences (Harbin, China). Enzyme activities were assessed using a soil enzyme activity test Kit (Suzhou Grace Biotechnology Co., Ltd., Suzhou, China). Specific measurement and analysis followed the manufacturer’s instructions using Multiskan sky (Thermo Fisher Scientific, Waltham. MA, USA). Assessment of the enzyme activities was as the following: soil urease (S-UE) was measured at 578 nm; neutral phosphatase (S-NP, G0306W) catalyzes p-nitrophenyl phosphate (pNPP) to produce a yellow product PNP, which has a maximum absorption peak at 405 nm, and the enzyme activity was measured by the rate of increase of PNP; solid polyphenol oxidase (S-PPO, G0311W) catalyzes gallic acid to produce gallium, which has a characteristic light absorption at 430 nm, reflecting polyphenol oxidase activity; solid sucrase (S-SC, G0302W) catalyzes the degradation of sucrose into reducing sugar and reacts with 3, 5-dinitrosalicylic acid to form colored metal amides with characteristic light absorption at 540 nm; soil catalase (S-CAT, G0303W) catalyzes hydrogen peroxide to produce water and oxygen and the remaining hydrogen peroxide reacts with a chromogenic probe to produce a colored substance with a maximum absorption at about 510 nm. Each treatment had three replicates and the experiment was conducted twice.

### Effects of soil extracts on alfalfa seedling and pathogen growth

To assess the effects of the autotoxic substances on alfalfa seed and fungal pathogens causing alfalfa root rot, soil extracts from the rhizosphere soil samples were prepared. Ten grams of soil were mixed with 250 mL distilled water and shaken for 24 h (Yang et al, 2009). The supernatant was filtered, distilled and concentrated to 1 mL/g of soil using a rotary evaporator (Strike 300, Guangzhou Wengdi Instruction Co., Ltd, China). The concentrated supernatants were filtered through a bacterial filter and stored at 4℃.

Alfalfa seeds were immersed in 1.5% sodium hypochlorite for 10 min, rinsed 5 times with distilled water, air dried, and soaked in the soil extracts (1 mL/g) for 30 min. Treated seeds were placed on sterile filter paper in a petri dish (50 seeds/dish). Two mL of soil extract was added to each dish and the dishes were incubated in a growth chamber at 25℃ with 12-h photoperiod. Alfalfa seeds treated with equal amount of distilled water were used as the control. Each treatment had three replicates and the experiment was conducted twice. Seed germination rate, root length and plant height of alfalfa were measured 7 days after incubation.

To assess the effects of soil extracts on pathogens causing alfalfa root rot, three fungal pathogens were used including *Fusarium trinctum* (MH894213), *F. acuminatum* (MK764994) and *F. oxysporum* (MK764964). The fungi were grown on potato dextrose agar (PDA) plates at 25°C for 5 days, and a mycelium plug (0.7 cm in diameter) was transferred onto PDA plates amended with the soil extract (final concentration V/V: 1%, 5% and 10%). PDA plates amended with equal amount of distilled water were used as controls. The plates were incubated at 26°C for 5 days and colony diameters were measured. Each treatment had three replicates and the experiment was conducted twice.

### Metabolomics analysis of alfalfa rhizosphere soil

The rhizosphere soil samples were extracted with methanol-water (v/v 3:1), ethyl acetate, L-2-chlorophenylalanine, and air dried. The air-dried soil extracts were dissolved in 20 μL methoxyamine salt and 30 μL BSTFA (containing 1% TMCS). Metabolites of the rhizosphere soils were assessed using 7890A gas chromatography-time-of-fight mass spectrometry (GC-TOF-MS) with DB-5MS capillary column (Agilent, USA) at Beijing Allwegene Co., Ltd. (Beijing, China). The Chroma TOF software (V 4.3x, LECO) was used to analyze the mass spectrum data, including peak extraction, baseline correction, deconvolution, peak integration and peak alignment (Kind *et al*., 2009). The LECO-Fiehn RTX5 database, including mass spectrometry, match and retention time index match, was used in the qualitative analysis of the substances.

### Validation of key metabolites

To determine the role of key metabolites in alfalfa CC, the effects of key metabolites from the rhizosphere soils on seed germination, seedling growth, and growth of root rot pathogens *F. trintum*, *F. acuminatum* and *F. oxysporum* were determined. The following experiments were conducted:

#### Effect of key metabolites on alfalfa seed and seedling

Alfalfa seeds disinfected as described above were immersed in different concentrations (10, 25, 50, 100 μg/mL) of key metabolites for 30 min. The seeds were placed in a sterile culture dish, covered with two layers of sterile filter paper, dripped with the corresponding concentration of key metabolites (1 mL/two days), and placed in a humidity chamber (>95% RH, 24℃ and 16/8 h light/dark). Each treatment had three replicates and the experiment was conducted twice. Seed germination rate, root length and plant height were measured 7 days after incubation.

#### Effect of key matabolites on pathogenic fungi causing alfalfa root rot

A mycelium plug (0.7 cm in diameter) of *F. trinctum*, *F. acuminatum* and *F. oxysporum* grown on PDA for 5 days was transferred onto PDA plates amended with different key metabolites at 10, 25, 50, and 100 μg/mL. The plates were incubated at 26°C in dark and colony diameters were measured 5 days after incubation. Plates amended with equal amount of sterile distilled water (SDW) were used as controls. Each treatment had three replicates and the experiment was conducted twice.

To evaluate effects of metabolites on spore germination, *F. trinctum*, *F. acuminatum* and *F. oxysporum* were grown on PDA at 25°C until colony diameters were 5 cm or larger. Conidia on the plates were washed with SDW, and the concentration was adjusted to 10^6^ spores/mL by counting using a hemocytometer. Different key metabolites were added into the conidial suspensions at concentrations of 10, 25, 50 and 100 μg/mL. Conidial suspensions amended with an equal volume of SDW served as a control. Spore suspensions were incubated at 25°C until spore germination rates were greater than 10%, and germinated spores were counted by counting 100 spores for each treatment in a replicate. To evaluate effects of metabolites on spore production, the three fungal cultures were grown as above. Different key metabolites at concentrations of 0, 10, 25, 50, or 100 μg/mL were added to each plate (20 mL/plate). Mycelia on the surface of the PDA plates were scraped off, and liquid on the plates was poured out after 20 min. The plates were then incubated at 26°C for 72 h, and 20 mL of SDW was added to a petri dish to wash the spores off the mycelium (Li et al, 2010). Spore suspension in a dish was collected in a tube (50 mL) and spore concentration was determined using a hemocytometer. Each treatment had three replicates and the experiment was conducted twice.

### Microbiological analysis of alfalfa rhizosphere soil

Total DNA was extracted from the rhizosphere soils using PowerSoil DNA Isolation Kit (MoBio Laboratories, CarIsbad, CA, USA). DNA concentration was quantified using a NanoDrop^TM^ 2000 spectrophotometer (Thermo Fisher Scientific, Waltham, MA, USA). Research on 16S rRNA/ITS sequencing and sequencing of the complete metagenomic sequence. The primer sets ITS1 (CTTGGTCATTTAGAGGAAGTAA) / ITS2 (TGCGTTCTTCATCGATGC) and 338F (ACTCCTACGGGAGGCAGCAG) / 806R (GGACTACHVGGGTWTCTAAT) were used to amplify target regions of fungal and bacterial genes, respectively.

The quality-checked DNA samples were then sequenced on Illumina Miseq PE300 platform. To guarantee the quality of data for downstream analysis, Trimmomatic was used to remove raw reads with tail end quality score < 20. Data pre-processing was conducted to obtain a good sequence. Through the sorting operation, the sequences were divided into different groups according to their similarity, and a group was an OTU. All sequences were divided into OTU according to different similarity level, and OTUs under 97% similarity level could be analyzed statistically (Edgar, 2013). The data were extracted based on the OTU clustering results, and the Alpha (Amato et al, 2013) and Beta (Jiang et al, 2013) analyses were carried out using Qiime (Version 1.8, http://qiime.org/), uclust (Version 1.2.22, http://www.drive5.com/uclust/downloads1_2_22q.html) and usearch (Version 10.0.240, http://www.drive5.com/usearch/)

### Validation of effects of p-coumaric acid in greenhouse

Alfalfa seeds were surface disinfested as described above and planted in a seedling tray in a greenhouse at 25±2°C. Seven-day-old seedlings were transplanted in pots (5 plants/per pot). Plants in 5 pots were treated with p-coumaric acid 3 days after transplanting at 10, 25, 50, or 100 µg/mL, respectively (1 mL/per plant). Treatment with p-coumaric acid was applied once every three days, and plants in 5 pots treated with SDW were used as controls. Alfalfa rhizosphere soils were collected as described above after treatment with p-coumaric acid for 5 times (i.e., 15 days after transplanting). Soil samples were stored at -80°C for microbiological analysis. Soil DNA extraction and microbiological analysis were as described above.

## Acknowledgements

The authors of this article would like to commemorate Prof. Pingsheng Ji from the Department of Plant Pathology at the University of Georgia. He is the co-author of this article. He passed away in May this year. This study was funded by National Natural Science Foundation of China (31971760), and the Heilongjiang Collaborative Innovation and Extension System of Modern Agricultural Industry Technology of Forage and Feed.

## Author contributions

RT performed all experiments. WY and JX provided help of the experiments. YG designed the study and modify the majority of the manuscript. PS provided comments on the manuscript. All authors read and approved the final manuscript.

